# Evidence for a hydrogen sulfide-sensing E3 ligase in yeast

**DOI:** 10.1101/2021.01.06.425657

**Authors:** Zane M. Johnson, Yun Wang, Benjamin M. Sutter, Benjamin P. Tu

## Abstract

In yeast, control of sulfur amino acid metabolism relies upon Met4, a transcription factor which activates the expression of a network of enzymes responsible for the biosynthesis of cysteine and methionine. In times of sulfur abundance, the activity of Met4 is repressed via ubiquitination by the SCF^Met30^ E3 ubiquitin ligase, but the mechanism by which the F-box protein Met30 senses sulfur status to tune its E3 ligase activity remains unresolved. Herein, we show that Met30 responds to flux through the transsulfuration pathway to regulate the *MET* gene transcriptional program. In particular, Met30 is responsive to the biological gas hydrogen sulfide, which is sufficient to induce ubiquitination of Met4 *in vivo*. Additionally, we identify important cysteine residues in Met30’s WD-40 repeat region that sense the availability of sulfur in the cell. Our findings reveal how SCF^Met30^ dynamically senses the flow of sulfur metabolites through the transsulfuration pathway to regulate synthesis of these special amino acids.

## INTRODUCTION

The biosynthesis of sulfur-containing amino acids supplies cells with increased levels of cysteine and methionine, as well as their downstream metabolites glutathione and S-adenosylmethionine (SAM). Glutathione serves as a redox buffer to maintain the reducing environment of the cell and provide protection against oxidative stress, while SAM serves as the methyl donor for nearly all methyltransferase enzymes (1,2). In the yeast *Saccharomyces cerevisiae*, biosynthesis of all sulfur metabolites can be performed *de novo* via enzymes encoded in the gene transcriptional network known as the *MET* regulon. Activation of the *MET* gene transcriptional program under conditions of sulfur starvation relies on the transcription factor Met4 and additional transcriptional co-activators that allow Met4 to be recruited to the *MET* genes (3,4).

When yeast cells sense sufficiently high levels of sulfur in the environment, the *MET* gene transcriptional program is negatively regulated by the activity of the SCF E3 ligase Met30 (SCF^Met30^) through ubiquitination of the master transcription factor Met4 (5). Met4 is unique as an E3 ligase substrate as it contains an internal ubiquitin interacting motif (UIM) which folds in and caps the growing ubiquitin chain generated by SCF^Met30^, resulting in a proteolytically stable but transcriptionally inactive oligo-ubiquitinated state (6). Upon sulfur starvation, SCF^Met30^ ceases to ubiquitinate Met4, allowing Met4 to become deubiquitinated and transcriptionally active.

Since its discovery, much effort has gone into understanding how Met30 senses the sulfur status of the cell. Several mechanisms have been attributed to Met30 to describe how Met4 and itself work together to regulate levels of *MET* gene transcripts in response to the availability of sulfur or the presence of toxic heavy metals (7). After the discovery that Met30 is an E3 ligase that negatively regulates Met4 through ubiquitin-dependent and both proteolysis-dependent and independent mechanisms (8-10), it was found that Met30 dissociates from SCF complexes upon cadmium addition, resulting in the disruption of the aforementioned ubiquitin-dependent regulatory mechanisms (11). It was later reported that this cadmium-specific dissociation of Met30 from SCF complexes is mediated by the Cdc48/p97 AAA+ ATPase complex, and that Met30 ubiquitination is required for Cdc48 to strip Met30 from these complexes (12). In parallel, attempts to identify the sulfur metabolic cue sensed by Met30 suggested that cysteine, or possibly some downstream metabolite, was required for the degradation of Met4 by SCF^Met30^, although glutathione was reportedly not involved in this mechanism (13,14). A genetic screen for mutants that fail to repress *MET* gene expression found that *cho2Δ* cells, which are defective in the synthesis of phosphatidylcholine (PC) from phosphatidylethanolamine (PE), results in elevated SAM levels and deficiency in cysteine levels, implying reduced flux through the transsulfuration pathway (15). However, while Met30 and Met4 have been studied extensively for over two decades, the biochemical mechanisms by which Met30 senses and responds to the presence or absence of sulfur remains incomplete (15).

Herein, we utilize prototrophic yeast strains grown in sulfur-rich and sulfur-free respiratory conditions to further elucidate the mechanism by which Met30 senses sulfur. Using genetic blockades in the sulfur metabolic pathway to identify the signal for sulfur sufficiency, we find that the flux through the transsulfuration pathway is key for Met30 sulfur sensing. The major enzymes responsible for transsulfuration, cystathionine β-synthase (Cys4) and cystathionine γ-lyase (Cys3), are known to be promiscuous in their substrate specificity and can produce biologically meaningful amounts of the gasotransmitter sulfide (H_2_S) as a side reaction (16). We find that yeast strains incapable of producing or consuming sulfide are still able to respond to low quantities of exogenous sulfide with respect to Met30 E3 ligase activity. These data, along with Met30 cysteine to serine point mutants defective in sulfur sensing, provide evidence that Met30 may directly sense the generation of sulfide gas as a signal of sulfur sufficiency using key cysteine residues in Met30’s WD-40 repeat region.

## RESULTS

### SYNTHESIS OF CYSTEINE IS MORE IMPORTANT THAN METHIONINE FOR MET4 UBIQUITINATION

Previous work in our lab has characterized the metabolic and cellular response of yeast cells following switch from rich lactate media (YPL) to minimal lactate media (SL) (17-23). Under such respiratory conditions, yeast cells engage regulatory mechanisms that might otherwise be subject to glucose repression. Among other phenotypes, this switch results in the acute depletion of sulfur metabolites and the activation of the *MET* gene regulon (18,23). To better study the response of yeast cells to sulfur starvation, we reformulated standard minimal, defined media to contain no sulfate, as prototrophic yeast can assimilate sulfur in the form of inorganic sulfate into reduced sulfur metabolites. After switching cells from YP lactate media (Rich) to the new minimal sulfur-free lactate media (−Sulfur), we found that Met30 and Met4 quickly responded to sulfur starvation through the extensively studied ubiquitin-dependent mechanisms regulating Met4 activity (Fig. 1A) (5,6,9,11,24). As previously observed, the deubiquitination of Met4 resulted in the activation of the *MET* genes (Fig. 1B) and corresponded well with changes in observed sulfur metabolite levels (Fig. 1C). Addition of sulfur metabolites quickly rescued Met30 activity and resulted in the re-ubiquitination of Met4 and the repression of the *MET* genes.

**Figure 1.**
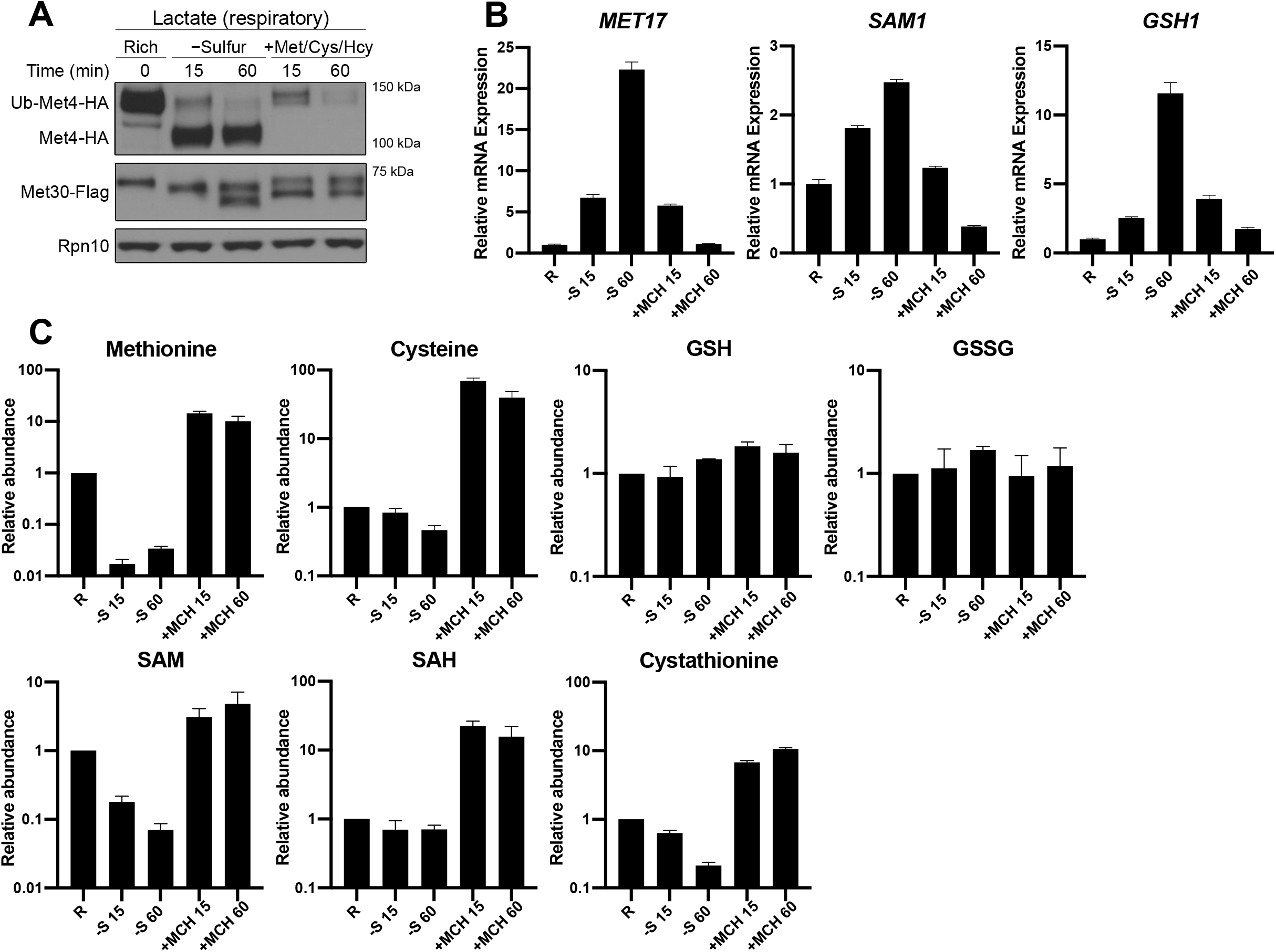
Met30 and Met4 response to sulfur starvation and repletion under respiratory growth conditions. (A) Western blot analysis of Met30 and Met4 over the sulfur starvation time course. Yeast containing endogenously tagged Met30 and Met4 were cultured in rich lactate media (Rich) overnight to mid log phase before switching cells to sulfur-free lactate media (−sulfur) for 1 h, followed by the addition of a mix of the sulfur containing metabolites methionine, homocysteine, and cysteine at 0.5 mM each (+Met/Cys/Hcy). Rpn10 is used as the loading control. (B) Expression of *MET* gene transcript levels was assessed by qPCR over the time course shown in (A). Data are presented as mean and SEM of technical triplicates. (C) Levels of key sulfur metabolites were measured over the same time course as in (A) and (B), as determined by LC-MS/MS. Data represent the mean and SD of two biological replicates.

As previously noted, Met4 activation in response to sulfur starvation results in the emergence of a second, faster-migrating proteoform of Met30, which disappears after rescuing yeast cells with sulfur metabolites (15). We found that the appearance of this proteoform is dependent on both *MET4* and new translation, as it was not observed in either *met4Δ* cells or cells treated with cycloheximide during sulfur starvation (Fig. S1A and C). Additionally, this proteoform persists after rescue with a sulfur source in the presence of a proteasome inhibitor (Fig. S1B).

We hypothesized that this faster-migrating proteoform of Met30 might be the result of translation initiation at an internal methionine residue. In support of this possibility, mutation of methionine residues 30, 35, and 36 to alanine blocked the appearance of a lower form during sulfur starvation (Fig. S1D). Conversely, deletion of the first 20 amino acids containing the first three methionine residues of Met30 resulted in expression of a Met30 proteoform that migrated at the apparent molecular weight of the wild type (WT) short form and did not generate a new, even-faster migrating proteoform under sulfur starvation (Fig. S1D). Moreover, the Met30^M30/35/36A^ and Met30^Δ1-20^ strains expressing either solely the long or short form of the Met30 protein had no obvious phenotype with respect to Met4 ubiquitination or growth in high or low sulfur media (Fig. S1E). We conclude that the faster-migrating proteoform of Met30 produced during sulfur starvation has no discernible effect on sulfur metabolic regulation under these conditions.

The sulfur amino acid biosynthetic pathway is bifurcated into two branches at the central metabolite homocysteine, where this precursor metabolite commits either to the production of cysteine or methionine (Fig. 2A). After confirming Met30 and Met4 were responding to sulfur starvation as expected, we sought to determine which sulfur metabolite is the signal for sulfur sufficiency for Met30. Deletion of *SAH1*, the enzyme responsible for recycling SAH back into homocysteine and adenosine after a successful SAM-dependent methylation reaction, results in the accumulation of SAH and competitive inhibition of SAM-dependent methyltransferases (25,26). Addition of SAM to this strain after starving cells for sulfur did not rescue Met4 ubiquitination status, while the addition of homocysteine was able to do so (Fig. 2B). Likewise, addition of methionine to the SAM-auxotrophic *sam1Δsam2Δ* double mutant did not rescue Met4 ubiquitination (Fig. 2B). These data suggest that no intermediate of the methionine/SAM branch of sulfur metabolism is the signal for sulfur sufficiency for Met30. To determine whether the synthesis of methionine is necessary to rescue Met30 activity, cells lacking methionine synthase (*met6Δ*) were fed either homocysteine or methionine after switching to sulfur-free lactate (−Sulfur) media. Interestingly, *met6Δ* cells fed homocysteine were still able to ubiquitinate and degrade Met4, while methionine-fed cells appeared to oligo-ubiquitinate and stabilize Met4 (Fig. 2C). These observations are consistent with previous reports and suggest Met30 and Met4 interpret sulfur sufficiency through both branches of sulfur metabolism to a degree (5,9,10,13-15), with the stability of Met4, but not the E3 ligase activity of Met30, apparently dependent on the methionine branch.

**Figure 2.**
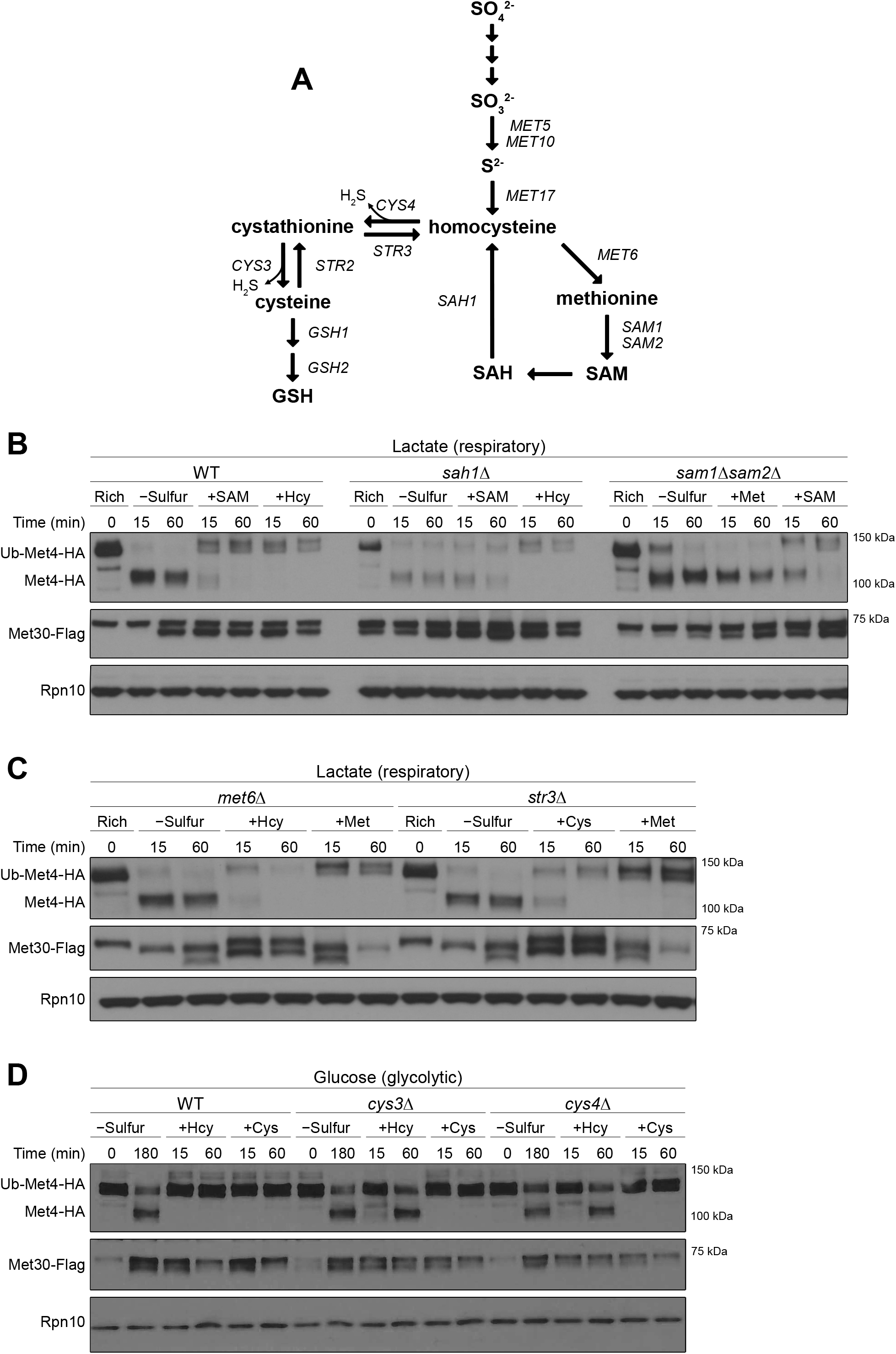
Synthesis of cysteine is more important than methionine for Met4 ubiquitination. (A) Simplified diagram of the yeast sulfur amino acid biosynthetic pathway. (B) Western blots of Met4 ubiquitination status in response to rescue with various sulfur metabolites in the methyl cycle in WT, *sah1Δ*, and *sam1Δ/sam2Δ* strains. Cells were grown in “Rich” YPL and switched to “−sulfur” SFL for 1 h to induce sulfur starvation before the addition of either 0.5 mM homocysteine (+Hcy), 0.5 mM methionine (+Met), or 0.5 mM *S*-adenosylmethionine (+SAM). (C) Western blots of Met4 ubiquitination status in *met6Δ* or *str3Δ* strains in response to rescue with various sulfur metabolites. Cells were grown in “Rich” YPL and switched to “−sulfur” SFL for 1 h to induce sulfur starvation before the addition of either 0.5 mM homocysteine (+Hcy), 0.5 mM methionine (+Met), or 0.5 mM cysteine (+Cys). (D) Western blots of Met4 ubiquitination status in WT, *cys3Δ* and *cys4Δ* strains produced in the S288C background in response to rescue with sulfur metabolites in the transsulfuration pathway. Cells were grown in sulfur free glucose media supplemented with methionine to log phase before removing methionine and switching to sulfur free glucose media for 3 h to induce sulfur starvation. Subsequently, either 0.5 mM homocysteine (+Hcy) or 0.5 mM cysteine (+Cys) was added to cells.

To determine whether Met30 specifically responds to the cysteine branch, cells lacking cystathionine β-lyase (*str3Δ*), the enzyme responsible for the conversion of cystathionine to homocysteine, were starved of sulfur and fed either cysteine or methionine. This mutant is incapable of synthesizing methionine from cysteine via the two-step conversion of cysteine into the common precursor metabolite homocysteine. Our results show cysteine was able to rescue Met30 activity even in a *str3Δ* mutant, further suggesting cysteine or a downstream metabolite, and not methionine, as the signal of sulfur sufficiency for Met30 (Fig. 2C). Together, these results suggest that the signal for the Met30-dependent re-ubiquitination of Met4 after sulfur starvation is independent of any sulfur metabolite within the methyl cycle itself, and that the ability of Met30 to sense sulfur-sufficiency in response to rescue with methionine or SAM is dependent on a fully functional methyl cycle capable of regenerating homocysteine.

Transsulfuration in the direction of cysteine synthesis requires the pyridoxal 5’-phosphate (PLP)-dependent enzymes cystathionine β-synthase (*CYS4*) and cystathionine γ-lyase (*CYS3*), and while these genes are essential in the CEN.PK background, they can be deleted in the S288C strain background. Although the *cys4Δ* and *cys3Δ* mutants grew poorly in media with lactate as a carbon source, we were able to test the ability of these mutants to sense homocysteine as a sulfur source in sulfur-free glucose media. Homocysteine addition to either *cys4Δ* or *cys3Δ* cells resulted in the transient re-ubiquitination of Met4 by Met30 at the 15 minute time point, but Met4 returned to its deubiquitinated state after 60 minutes. Notably, cysteine, which is downstream of both Cys4p and Cys3p, fully rescued the E3 ligase activity of Met30 in mutants lacking either of these enzymes (Fig. 2D).

### HYDROGEN SULFIDE IS A SIGNAL FOR SULFUR SUFFICIENCY

The unusual result of the homocysteine rescue in the *cys4Δ* and *cys3Δ* mutants led us to speculate that it is something unique about these enzymes that is contributing to the Met30 sulfur sensing mechanism. It is known that these two enzymes produce the biologically relevant gas hydrogen sulfide (H_2_S), an important signaling molecule responsible for a wide range of biological effects (27,28). We suspected that our transient homocysteine rescue may be attributable to the promiscuous nature of the substrate specificity of these enzymes, which can produce hydrogen sulfide from cysteine and homocysteine as a byproduct (16). To test if Met30 can sense hydrogen sulfide, we deleted both *MET10* and *MET17* and treated the sulfur-starved yeast with 20 µM disodium sulfide. Deletion of *MET10*, one of the two genes encoding the heterotetrameric enzyme sulfite reductase, as well as *MET17*, which encodes homocysteine synthase - results in a yeast strain that cannot produce sulfide from inorganic sulfate via sulfate assimilation, and even when provided sulfide exogenously, cannot incorporate the sulfide into organic sulfur metabolites. Remarkably, treatment of the *met10Δmet17Δ* double mutant with disodium sulfide was able to rescue Met30 E3 ligase activity to the degree seen in WT yeast (Fig. 3A). Measurement of sulfur-containing metabolites over the time course confirmed that the double mutant was not incorporating sulfide into homocysteine or cystathionine, suggesting that low quantities of sulfide alone are sufficient for Met30 sulfur sensing (Fig. 3B).

**Figure 3.**
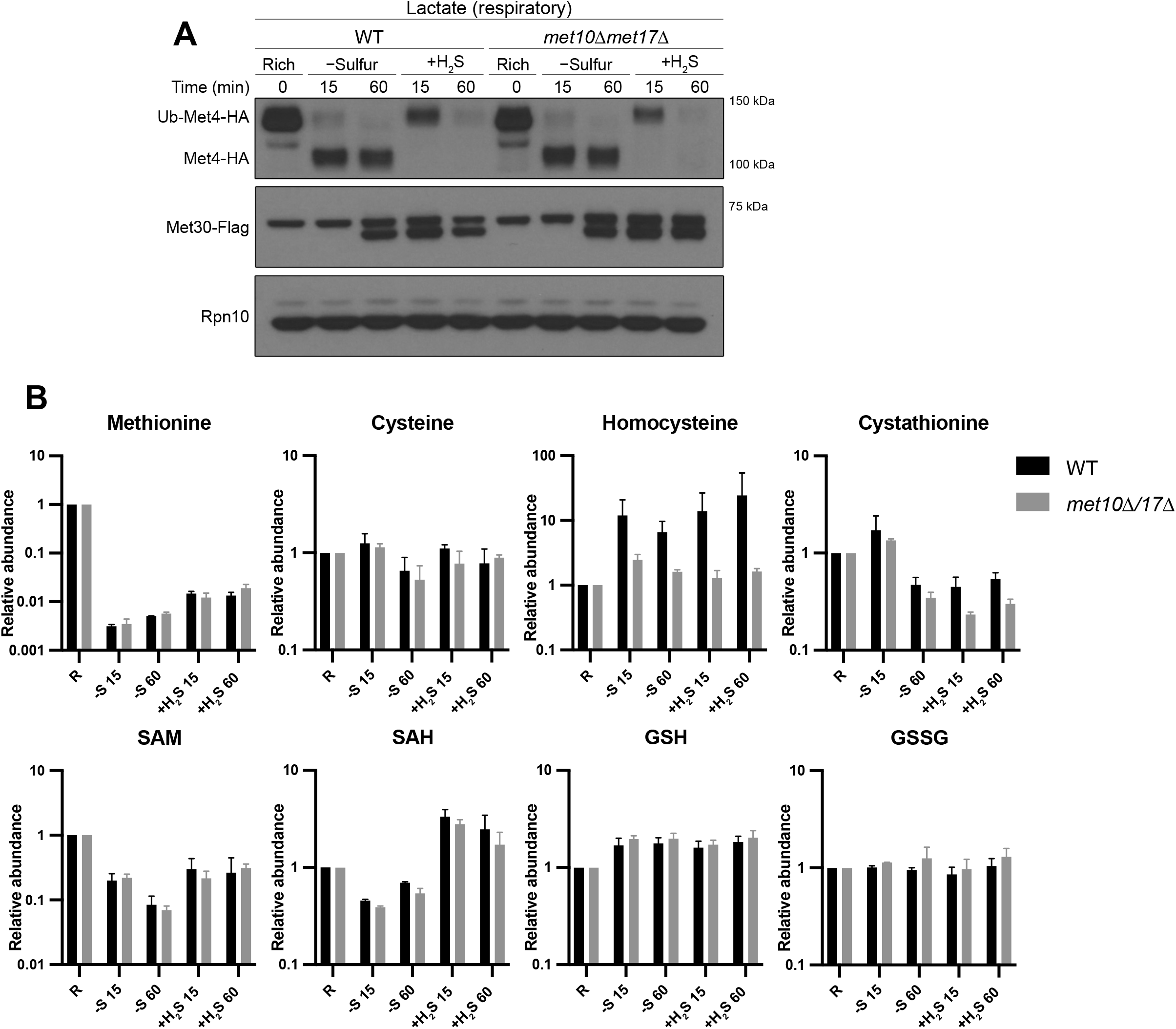
Hydrogen sulfide is a signal for sulfur sufficiency. (A) Western blot analysis of Met30 and Met4 ubiquitination in WT and *met10Δ/met17Δ* yeast grown in “Rich” YPL and switched to “−sulfur” SFL for 1 h to induce sulfur starvation before the addition of 20 µM disodium sulfide. Note that the *met10Δ/met17Δ* strain cannot produce sulfide from inorganic sulfate or utilize exogenous sulfide to produce homocysteine. (B) Levels of key sulfur metabolites were measured over the same time course as in (A), as determined by LC-MS/MS. Data represent the mean and SD of two biological replicates.

### MET30 CYSTEINE POINT MUTANTS EXHIBIT DYSREGULATED SULFUR SENSING *IN VIVO*

The synthesis of cysteine from homocysteine contributes to the production of the downstream tripeptide glutathione (GSH), which exists at millimolar concentrations in cells and is the major cellular reductant for buffering against oxidative stress (29,30). Specifically, glutathione serves to neutralize reactive oxygen species such as peroxides and free radicals, detoxify heavy metals, and preserve the reduced state of protein thiols (31,32). Like glutathione, sulfide is also nucleophilic and reductive by nature – and may directly reduce protein disulfides or form persulfides on cysteine residues (33). Considering the relatively high number of cysteine residues in Met30 (Fig. 4A) — as well as Met30’s apparent ability to successfully re-ubiquitinate Met4 in response to treatment with the membrane-permeable reducing agent DTT (Fig. S2A, third lane) — we sought to determine whether specific cysteine residues play key roles in the sensing mechanism. Through site-directed mutagenesis of Met30 cysteines individually and in clusters (Fig. 4A and 4B), we observed that mutation of cysteines in the WD-40 repeat regions of Met30 with the highest concentration of cysteine residues (WD-40 repeat regions 4 and 8) resulted in dysregulated Met4 ubiquitination status (Fig. 4B) and *MET* gene expression (Fig. 4C). Specifically, conservatively mutating these cysteines to serine residues mimics the reduced state of the Met30 protein, resulting in constitutive ubiquitination of Met4 by Met30 even when cells are starved of sulfur. The mixed population of ubiquitinated and deubiquitinated Met4 in the mutant strains resulted in reduced induction of *SAM1* and *GSH1*, while *MET17* appears to be upregulated in the mutants but is largely insensitive to the changes in the sulfur status of the cell. Interestingly, a single cysteine to serine mutant, C414S, phenocopies the grouped cysteine to serine mutants C414/426/436/439S (data not shown) and C614/616/622/630S. These mutants also exhibit slight growth phenotypes when cultured in both rich and −sulfur lactate media supplemented with homocysteine (Fig. 4D). Furthermore, these point mutants only affect Met4 ubiquitination in the context of sulfur starvation, as strains expressing these mutants exhibited a normal response to cadmium as evidenced by rapid deubiquitination of Met4 (Fig. S2B).

**Figure 4.**
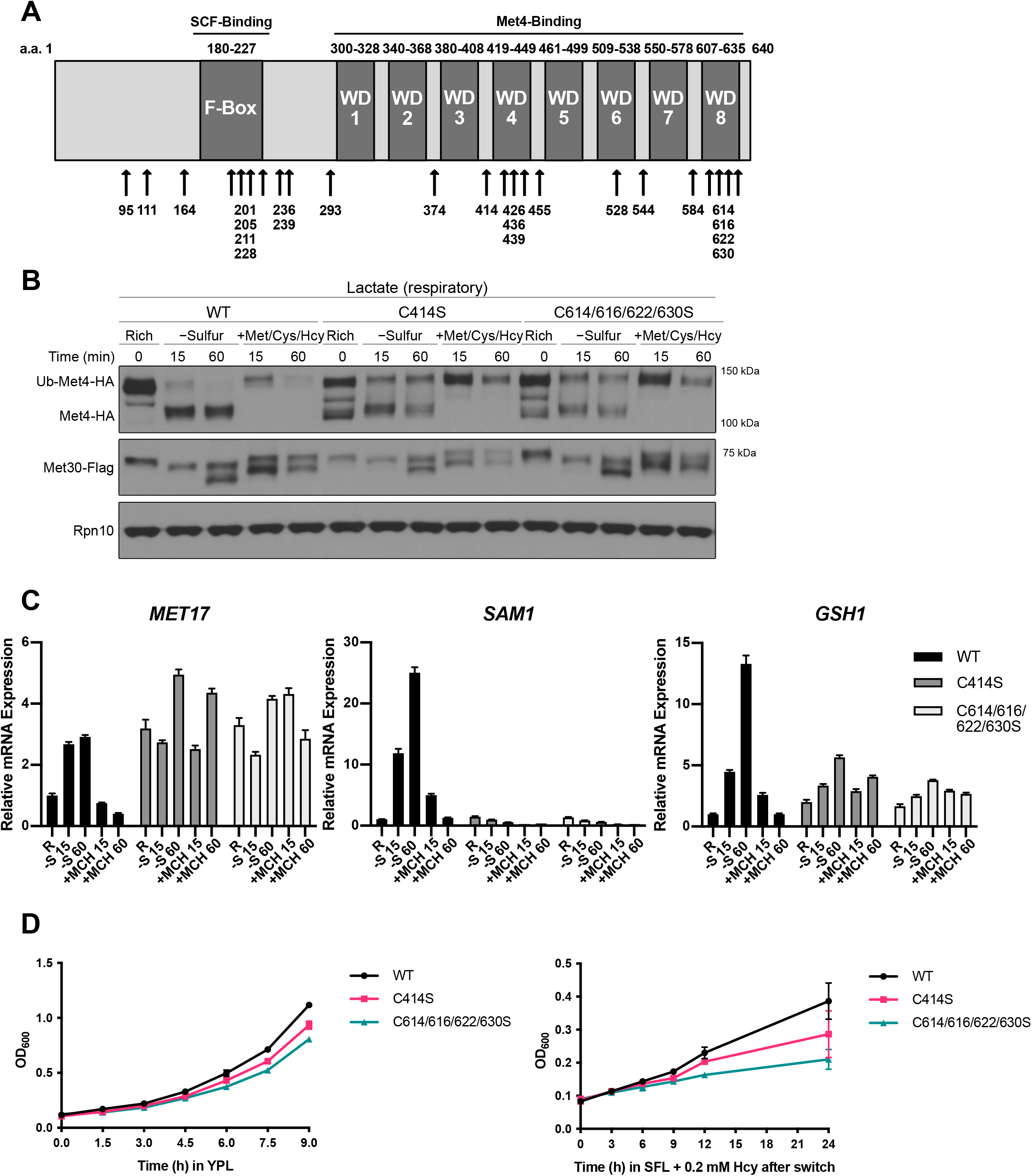
Met30 cysteine point mutants display dysregulated sulfur sensing. (A) Schematic of Met30 protein architecture and cysteine residue location. (B) Western blot analysis of Met30 and Met4 ubiquitination status in WT and two cysteine to serine mutants, C414S and C614/616/622/630S. (C) *MET* gene transcript levels over the same time course as (A) for the three strains, as assessed by qPCR. Data are presented as mean and SEM of technical triplicates. (D) Growth curves of the three yeast strains used in (A) and (B) in sulfur-rich YPL media or −sulfur SFL media supplemented with 0.2 mM homocysteine. Cells were grown to mid-log phase in YPL media before pelleting, washing with water, and back-diluting yeast into the two media conditions. Data represent mean and SD of technical triplicates.

Given the requirement of key cysteine residues within Met30 for sulfur sensing, we next sought to determine whether Met30 might undergo reversible redox modifications in response to sulfur. Moreover, sulfide is redox-active and can induce the persulfidation of cysteine residues in proteins. We performed the sulfur starvation and sulfide rescue time course, rapidly quenched cells with TCA to preserve the oxidation status of cysteine residues, and then subjected denatured Met30 to alkylation by the thiol-specific reagent mPEG2K-mal (methoxy-polyethylene glycol maleimide, average MW = 2 kDa), which is specific for the reduced form of cysteine. Met30 migrated at an apparent molecular weight of ∼150 kDa, consistent with modification of the majority of its 23 cysteine residues, and did not exhibit any significant mass shift on Western blots in response to sulfur starvation using this method (Fig. S3A).

In addition, we utilized a second strategy to detect oxidation by initially alkylating any reduced cysteine residues in Met30 with NEM immediately following lysis. Met30 was then treated with the non-thiol based reducing agent TCEP to reduce any oxidized cysteine residues. We then added the mPEG2K-mal reagent to modify any newly reduced Met30 cysteine residues (Fig. S3B). However, the data showed no observable retardation in the mobility of Met30 in response to sulfur starvation that would be suggestive of oxidative cysteine modification. Taken together, these two approaches suggest that nearly all of the 23 cysteine residues in Met30 are in the reduced form in both sulfur-replete and sulfur-deficient conditions. Although we show that key cysteine residues in Met30 are clearly involved in sulfur-sensing, these thiol-trapping strategies did not reveal evidence of sulfide-mediated changes to the modification of these cysteine residues by alkylating agents (see Discussion).

## DISCUSSION

The unique chemistry offered by sulfur and sulfur-containing metabolites renders many of the biochemical reactions required for life possible. The ability to carefully regulate the levels of these sulfur-containing metabolites is of critical importance to cells as evidenced by an exquisite sulfur-sparing response. Sulfur starvation induces the transcription of *MET* genes and specific isozymes, which themselves contain few methionine and cysteine residues (34). Furthermore, along with the dedicated cell cycle F-box protein Cdc4, Met30 is the only other essential F-box protein in yeast, linking sulfur metabolite levels to cell cycle progression (35,36). Our findings provide evidence that the key sensor of sulfur levels in yeast, Met30, may directly sense sulfur in the form of sulfide via key cysteine residues in its WD-40 repeat region. While much work has been done to characterize the molecular basis of sulfur metabolic regulation in yeast between Met30 and Met4, this work describes a biochemical basis for sulfur sensing by the Met30 E3 ligase (Fig. 5).

**Figure 5.**
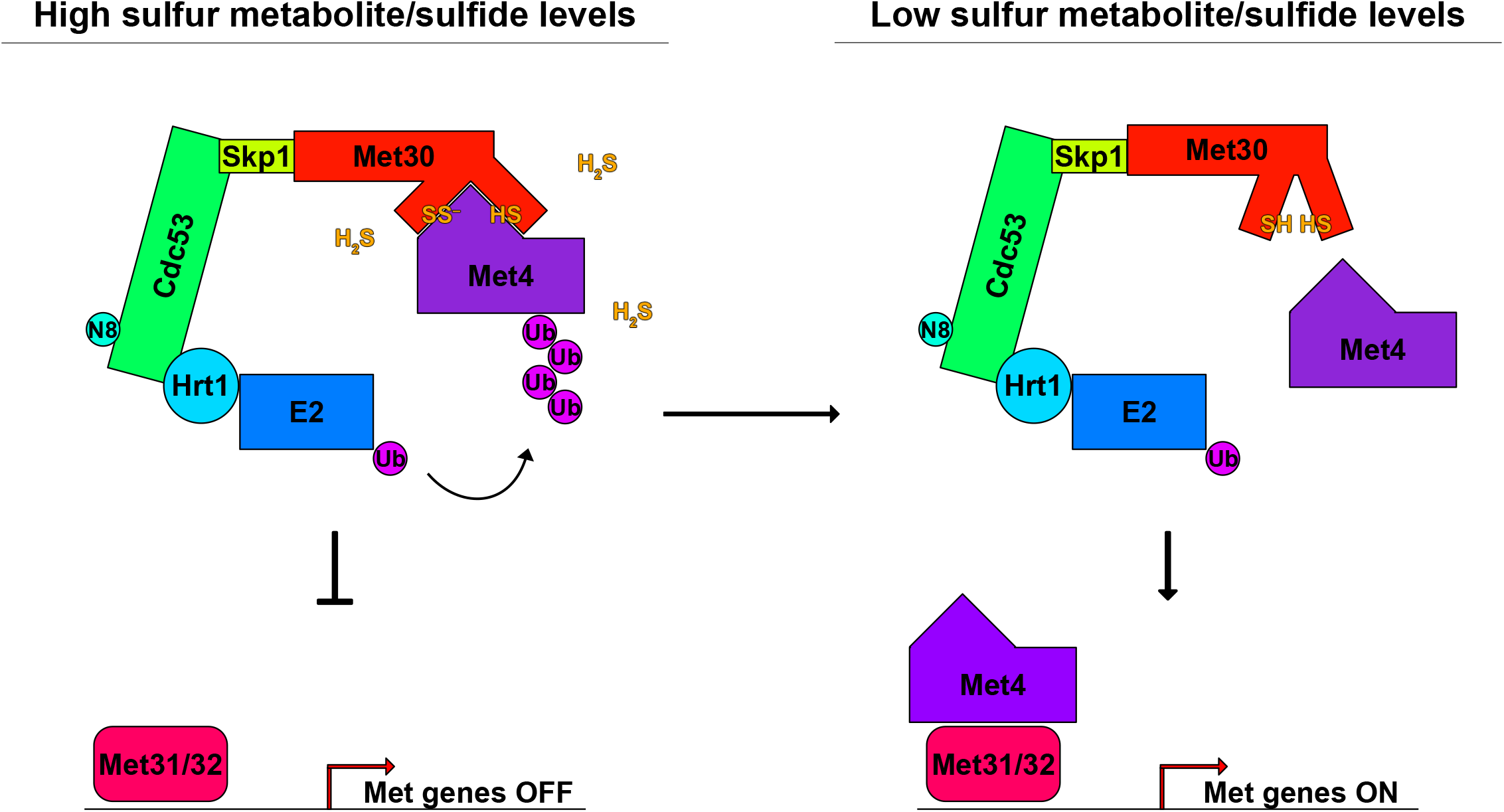
Model for sulfur-sensing and *MET* gene regulation by the SCF^Met30^ E3 ligase. In conditions of high sulfur metabolite and sulfide levels, cysteine residues in the WD-40 repeat region of Met30 are reduced or potentially persulfidated, allowing Met30 to bind and facilitate ubiquitination of Met4 in order to inhibit the transcriptional activation of the *MET* regulon. Upon sulfur starvation, Met30 releases Met4 to be deubiquitinated and activate the *MET* gene transcriptional program.

Of the small family of biologically active gases known as gasotransmitters, sulfide might be the most unusual. Notoriously toxic at high levels, the beneficial effects of low levels of sulfide exposure have been known for over half a century (37). Interest into the effects of the signaling molecule were piqued in 2005, when it was reported that exposure of mice to low levels of the gas (20-80ppm) resulted in a rapid decrease in energy expenditure and induced an apparent “suspended animation-like state” that, incredibly, was shown to be completely reversible (38). Since then, the beneficial and cytoprotective effects of sulfide have been demonstrated in multiple tissues and cell types (39-41). The effect does not seem to be limited to multicellular eukaryotes. Indeed, it has recently been reported that treatment of yeast with sulfide increases their chronological lifespan (42).

While there is still debate concerning the prevalence and specificity of protein persulfides in cells, it is known that sulfide signaling through protein persulfidation can occur for several proteins (33). However, it is unclear if many of these reported persulfidated proteins are true physiological sensors of sulfide, or if they are simply victims of the reactive chemical species and any downstream effect is unintentional. Based on our findings, we propose that Met30 constitutes a physiological sensor of sulfide. Multiple considerations suggest why SCF^Met30^ might function as a bona fide sulfide-sensing E3 ligase. In response to sulfur starvation, yeast cells boost levels of enzymes involved in sulfate assimilation, the expensive process of reducing sulfur in the form of inorganic sulfate into sulfide, so that it may be incorporated into the carbon backbone of homoserine to produce homocysteine. This pathway is tightly regulated via the Met4 transcriptional program, as evidenced by the near-zero levels of sulfate assimilation enzyme activity in a *met4Δ* cell lysate (43). Sulfide sensing by Met30 has the implication that flux through sulfate assimilation and/or transsulfuration can rapidly report the levels and activity of these enzymes without requiring true end-product feedback inhibition of the Met4 transcriptional program. A sulfide-based signaling mechanism would also be readily reversible, require no new RNA or protein synthesis, and sulfide could be retrieved for synthesis of homocysteine. However, due to the transient and labile nature of cysteine persulfide (Cys-SSH) modifications, more sophisticated biochemical and structural methods are needed to elucidate how SCF^Met30^ senses sulfide. It is also possible that SCF^Met30^ could utilize a cysteine-coordinated metal ion that is responsive to sulfide.

The unusual and fleeting sulfide-based signal may also explain why the mechanism by which Met30 senses sulfur has been so elusive, as well as reconcile some previous observations. For example, the observation by Hansen and Johannesen that *MET* gene repression can be achieved with 2 mM methionine, but that 10 mM cysteine (with or without 2 mM DTT) is necessary for similar levels of repression, can be explained by the methionine-induced production of sulfide via transsulfuration, while ∼5-fold higher levels of cysteine may be required to produce similar levels of sulfide. Intriguingly, we found that DTT alone can result in Met4 ubiquitination (Fig. S2A, third lane). We speculate that addition of the potent reducing agent may liberate the sulfane sulfur of persulfides or polysulfides in cells, resulting in increased sulfide levels. A surprising relationship between sulfur and lipid metabolism was discovered in 2014 when it was reported that loss of *CHO2*, a gene encoding a methyltransferase involved in synthesizing phosphatidylcholine (PC) from phosphatidylethanolamine (PE), results in a strain that fails to repress the *MET* gene transcriptional program when sulfur levels are high. The *cho2Δ* strain is characterized by the accumulation of high levels of the methyl donor SAM, as well as reduced capacity for the synthesis of cysteine from homocysteine (13,22). The constitutive activation of the Met4 transcriptional program in the *cho2Δ* strain is consistent with reduced flux through the transsulfuration pathway and corresponding low levels of sulfide production via the transsulfuration enzymes Cys4p and Cys3p.

It is also worth noting the curious spacing and clustering of cysteine residues in Met30, with the highest density and closest spacing of cysteines found in two WD-40 repeats that are expected to be directly across from each other in the 3D structure (Figure 4A). That the mutation of these cysteine clusters to serine have the largest *in vivo* effect, but mutation of any one cysteine to serine (with the notable exception of Cys414) has no effect, implies some built-in redundancy in the cysteine-based, sulfide-sensing mechanism (Figure S2A). We speculate that persulfidation and/or sustained reduction of key Met30 cysteines permits binding to Met4, while low levels of sulfide produce biochemical and structural changes in the WD-40 repeat region that render Cys414 and/or other cysteine residues incapable of interaction with Met4. Future structural characterization of Met30 in its reduced, oxidized, or persulfidated states will be required to understand how Met30 uses sulfide to regulate its interaction with, and direct its E3 ligase towards, Met4. Nevertheless, our findings provide evidence that the master sensor of sulfur in yeast, Met30, uses gaseous sulfide as a signal for sulfur sufficiency.

## EXPERIMENTAL PROCEDURES

### Yeast strains, construction, and growth media

The prototrophic CEN.PK strain background (44) was used in all experiments with the exception of Figure 2D, where the S288C background was required to obtain the *cys4Δ* and *cys3Δ* mutants. Strains used in this study are listed in Table S1. Gene deletions were carried out using either tetrad dissection or standard PCR-based strategies to amplify resistance cassettes with appropriate flanking sequences, and replacing the target gene by homologous recombination (45). C-terminal epitope tagged strains were similarly made with the PCR-based method to amplify resistance cassettes with flanking sequences. Point mutations were made by cloning the gene into the tagging plasmids, making the specific point mutation(s) by PCR, and amplifying and transforming the entire gene locus and resistance markers with appropriate flanking sequences using the lithium acetate method.

Media used in this study: YPL (1% yeast extract, 2% peptone and 2% lactate); sulfur-free glucose and lactate media (SFD/L) media composition is detailed in Table S2, with glucose or lactate diluted to 2% each; YPD (1% yeast extract, 2% peptone and 2% glucose).

### Whole cell lysate Western blot preparation

Five OD_600_ units of yeast culture were quenched in 15% TCA for 15 min, pelleted, washed with 100% EtOH, and stored at −20°C. Cell pellets were resuspended in 325 μL EtOH containing 1 mM PMSF and lysed by bead beating. The lysate was separated from beads by inverting the screwcap tubes, puncturing the bottom with a 23G needle, and spinning the lysate at 2,500xg into an Eppendorf for 1 min. Beads were washed with 200 μL of EtOH and spun again before discarding the bead-containing screwcap tube and pelleting protein extract at 21,000xg for 10 min in the new Eppendorf tube. The EtOH was aspirated and EtOH precipitated protein pellets were resuspended in 150 μL of sample buffer (200 mM Tris pH 6.8, 4% SDS, 20% glycerol, 0.2 mg/ml bromophenol blue), heated at 42°C for 45 min, and debris was pelleted at 16,000xg for 3 min. DTT was added to a final concentration of 25 mM and incubated at RT for 30 min before equivalent amounts of protein were loaded onto NuPAGE 4-12% bis-tris or 3-8% tris-acetate gels. For the modified tag-switch protocol in Figure S3, cells were quenched in 20% TCA, pelleted, washed twice with ice cold acetone, air dried and stored at −20°C. Cell pellets were resuspended and bead beaten in 250 μL urea buffer (6 M urea, 1% SDS, 50 mM Tris-HCl pH 6.8, 1 mM EDTA, 2x Roche protease inhibitors, 1 mM PMSF, 10 μM leupeptin, 5 μM pepstatin A, 1 mM sodium orthovanadate) supplemented with 20 mM NEM and lysate was incubated at 42°C for 45 min. Lysate was pelleted at 21,000xg for 15 min, the supernatant moved to a new tube, and protein was TCA precipitated (20% FC) for 30 min on ice. Precipitated protein was pelleted at 21,000xg for 10 min, supernatant was aspirated, and the pellet was washed twice with ice cold acetone and air dried before resuspending in 250 μL urea buffer containing 10 mM TCEP. Resuspended protein samples were treated with 5 mM mPEG2K-mal at 42°C for 45 min before mixing with sample buffer, heating at 65°C for 10 min, and loading onto 3-8% tris-acetate gels.

### Western blots

Western blots were carried out by transferring whole cell lysate extracts or *in vitro* ubiquitination or binding assay samples onto 0.45 micron nitrocellulose membranes and wet transfers were carried out at 300 mA constant for 90 min at 4°C. Membranes were incubated with ponceau S, washed with TBST, blocked with 5% milk in TBST for 1 h, and incubated with 1:5000 Mouse anti-FLAG M2 antibody (Sigma, Cat#F3165), 1:5000 Mouse anti-HA(12CA5) (Roche, Ref#11583816001), 1:50,000 Rabbit anti-RPN10 (Abcam, ab98843), or 1:3000 Goat anti-Cdc53 (Santa Cruz, yC-17) in 5% milk in TBST overnight at 4°C. After discarding primary antibody, membranes were washed 3 times for 5 min each before incubation with appropriate HRP-conjugated secondary antibody for 1 h in 5% milk/TBST. Membranes were then washed 3 times for 5 min each before incubating with Pierce ECL western blotting substrate and exposing to film.

### RNA Extraction and Real Time Quantitative PCR (RT-qPCR) Analysis

RNA isolation of five OD_600_ units of cells under different growth conditions was carried out following the manufacture manual using MasterPure yeast RNA purification kit (epicentre). RNA concentration was determined by absorption spectrometer. 5 μg RNA was reverse transcribed to cDNA using Superscript III Reverse Transcriptase from Invitrogen. cDNA was diluted 1:100 and real-time PCR was performed in triplicate with iQ SYBR Green Supermix from BioRad.

Transcripts levels of genes were normalized to ACT1. All the primers used in RT-qPCR have efficiency close to 100%, and their sequences are listed below.

ACT1_RT_F TCCGGTGATGGTGTTACTCA

ACT1_RT_R GGCCAAATCGATTCTCAAAA

MET17_RT_F CGGTTTCGGTGGTGTCTTAT

MET17_RT_R CAACAACTTGAGCACCAGAAAG

GSH1_RT_F CACCGATGTGGAAACTGAAGA

GSH1_RT_R GGCATAGGATTGGCGTAACA

SAM1_RT_F CAGAGGGTTTGCCTTTGACTA

SAM1_RT_R CTGGTCTCAACCACGCTAAA

### Metabolite extraction and quantitation

Intracellular metabolites were extracted from yeast using a previous established method (46). Briefly, at each time point, ∼12.5 OD_600_ units of cells were rapidly quenched to stop metabolism by addition into 37.5 mL quenching buffer containing 60% methanol and 10 mM Tricine, pH 7.4. After holding at -40°C for at least 3 min, cells were spun at 5,000xg for 2 min at 0°C, washed with 1 mL of the same buffer, and then resuspended in 1 mL extraction buffer containing 75% ethanol and 0.1% formic acid. Intracellular metabolites were extracted by incubating at 75°C for 3 min, followed by incubation at 4°C for 5 min. Samples were spun at 20,000xg for 1 min to pellet cell debris, and 0.9 mL of the supernatant was transferred to a new tube. After a second spin at 20,000xg for 10 min, 0.8 mL of the supernatant was transferred to a new tube. Metabolites in the extraction buffer were dried using SpeedVac and stored at −80°C until analysis. Methionine, SAM, SAH, cysteine, GSH and other cellular metabolites were quantitated by LC-MS/MS with a triple quadrupole mass spectrometer (3200 QTRAP, AB SCIEX) using previously established methods (46). Briefly, metabolites were separated chromatographically on a C18-based column with polar embedded groups (Synergi Fusion-RP, 150 × 2.0 mm, 4 µ, Phenomenex), using a Shimadzu Prominence LC20/SIL-20AC HPLC-autosampler coupled to the mass spectrometer. Flow rate was 0.5 ml/min using the following method: Buffer A: 99.9% H2O/0.1% formic acid, Buffer B: 99.9% methanol /0.1% formic acid. T = 0 min, 0% B; T = 4 min, 0% B; T = 11 min, 50% B; T = 13 min, 100% B; T = 15 min, 100% B, T = 16 min, 0% B; T = 20 min, stop. For each metabolite, a 1 mM standard solution was infused into a Applied Biosystems 3200 QTRAP triple quadrupole-linear ion trap mass spectrometer for quantitative optimization detection of daughter ions upon collision-induced fragmentation of the parent ion [multiple reaction monitoring (MRM)]. The parent ion mass was scanned for first in positive mode (usually MW + 1). For each metabolite, the optimized parameters for quantitation of the two most abundant daughter ions (i.e., two MRMs per metabolite) were selected for inclusion in further method development. For running samples, dried extracts (typically 12.5 OD units) were resuspended in 150 mL 0.1% formic acid, spun at 21,000xg for 5 min at 4°C, and 125 μL was moved to a fresh Eppendorf. The 125 μL was spun again at 21,000xg for 5 min at 4°C, and 100 μL was moved to mass-spec vials for injection (typically 50 μL injection volume). The retention time for each MRM peak was compared to an appropriate standard. The area under each peak was then quantitated by using Analyst® 1.6.3, and were re-inspected for accuracy. Normalization was done by normalizing total spectral counts of a given metabolite by OD_600_ units of the sample. Data represents the average of two biological replicates.

## DATA AVAILABILITY

All data are included in the manuscript.

## SUPPORTING INFORMATION

This article contains supporting information.

## ACKNOWLEDGMENTS

We thank members of the Tu lab, Deepak Nijhawan, Hongtao Yu, and George DeMartino for helpful discussions.

## AUTHOR CONTRIBUTIONS

This study was conceived by Z.J. and B.P.T. B.M.S. performed Met30 cysteine point mutant strain construction, Y.W. performed cysteine point mutant cloning and protein purification, and all remaining experiments were directed and performed by Z.J. The paper was written by Z.J. and B.P.T. and has been approved by all authors.

## FUNDING AND ADDITIONAL INFORMATION

This work was supported by NIH grant R35GM136370, the Welch Foundation I-1797, and an HHMI-Simons Faculty Scholars Award to B.P.T. B.P.T. is an Investigator of the Howard Hughes Medical Institute.

## SUPPLEMENTAL FIGURE LEGENDS

**Figure S1.**
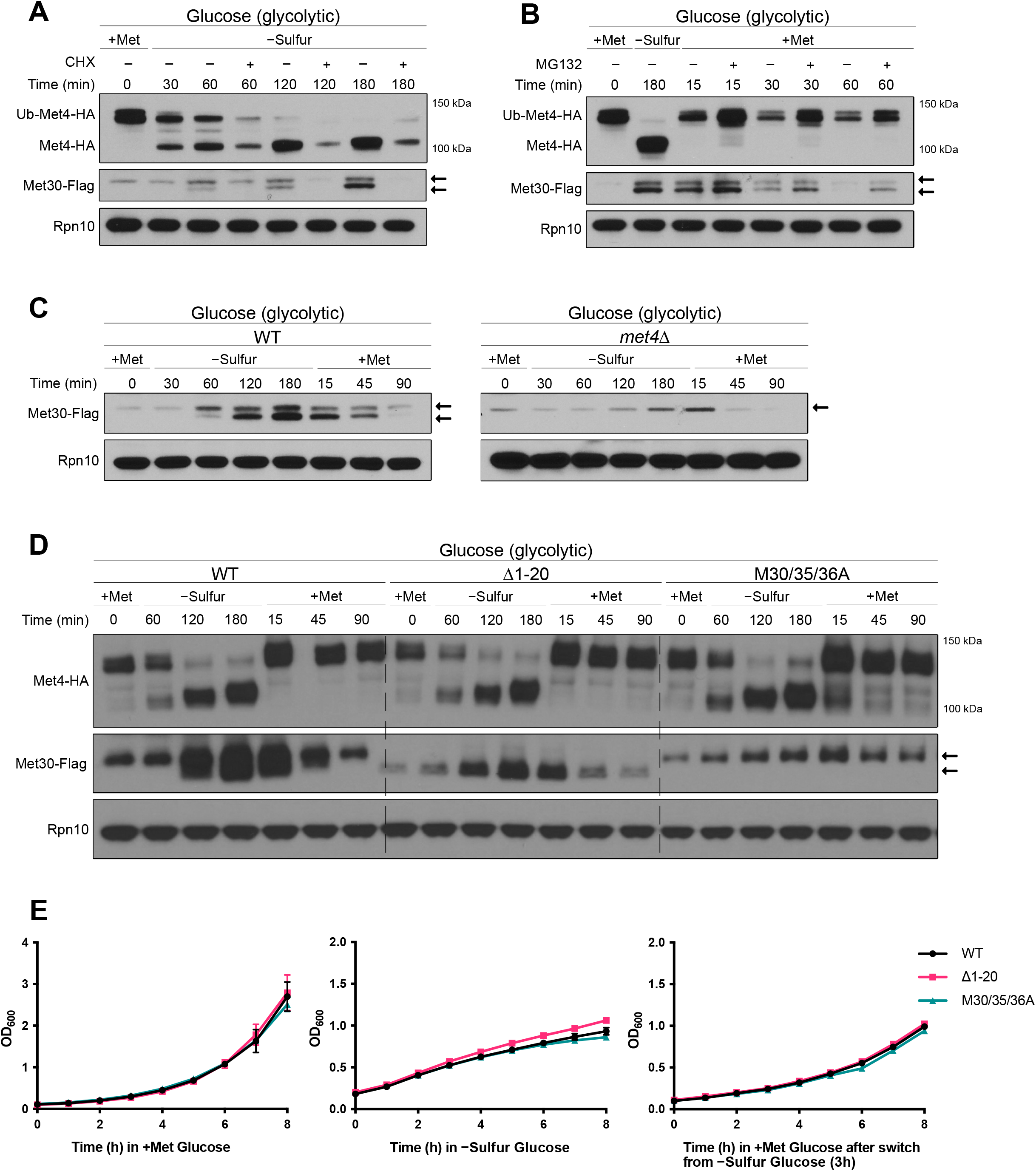
Characterization of the faster-migrating proteoform of Met30. (A) Western blot of yeast treated with 200 μg/ml cycloheximide during sulfur starvation demonstrates that production of the faster-migrating proteoform is dependent on new translation. (B) The faster-migrating proteoform persists after rescue from sulfur starvation when treated with a proteasome inhibitor. Cells were starved of sulfur for 3 h to accumulate the faster-migrating proteoform, and then sulfur metabolites were added back concomitantly with MG132 (50 μM). (C) The faster-migrating proteoform of Met30 is dependent on Met4. The *met4Δ* yeast strain does not produce the second proteoform of Met30 when starved of sulfur. (D) Western blot analysis of strains expressing either wild type Met30, Met30 Δ1-20aa, or Met30 M30/35/36A. Yeast cells harboring the N-terminal deletion of the first twenty amino acids of Met30 (which contain the first three methionine residues) or have the subsequent three methionine residues (M30/35/36) mutated to alanine do not create faster-migrating proteoforms. (E) Met30(Δ1-20aa) and Met30(M30/35/36A) strains do not exhibit any growth phenotypes in −sulfur glucose media with or without supplemented methionine. There are also no defects in growth rate following repletion of methionine. Data represent mean and SD of technical triplicates.

**Figure S2.**
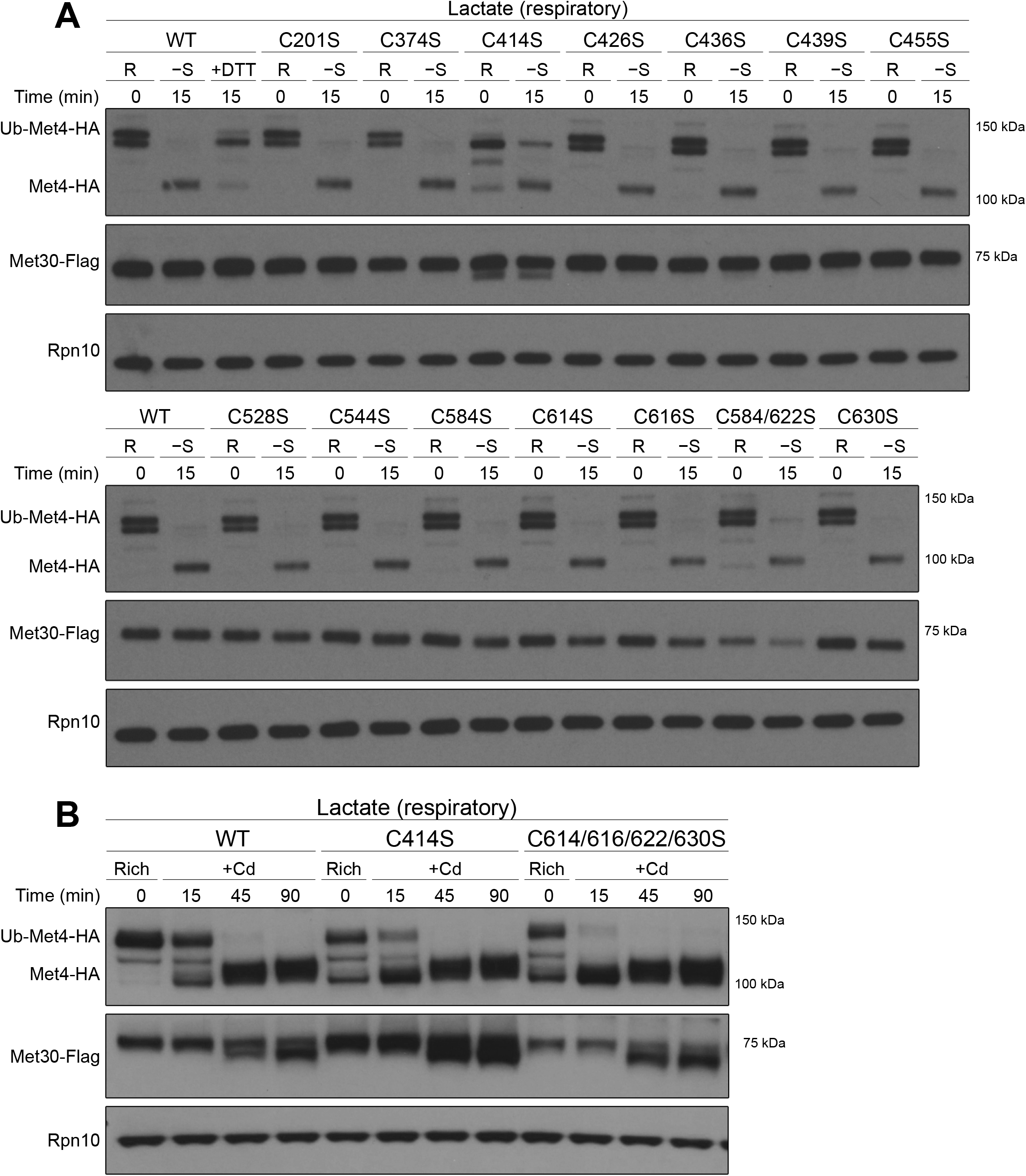
Identification of key cysteine residues in Met30 involved specifically in sulfur amino acid sensing. (A) Western blot analysis of Met4 ubiquitination in WT and various Met30 cysteine point mutants in rich and −sulfur media. Lane 3 depicts Met4 re-ubiquitination in response to treatment of sulfur-starved yeast cells with 5 mM DTT for 15 min. (B) Western blot analysis of Met30 and Met4 ubiquitination status in WT and two cysteine to serine mutants, C414S and C614/616/622/630S, following treatment with 500 µM CdCl_2_.

**Figure S3.**
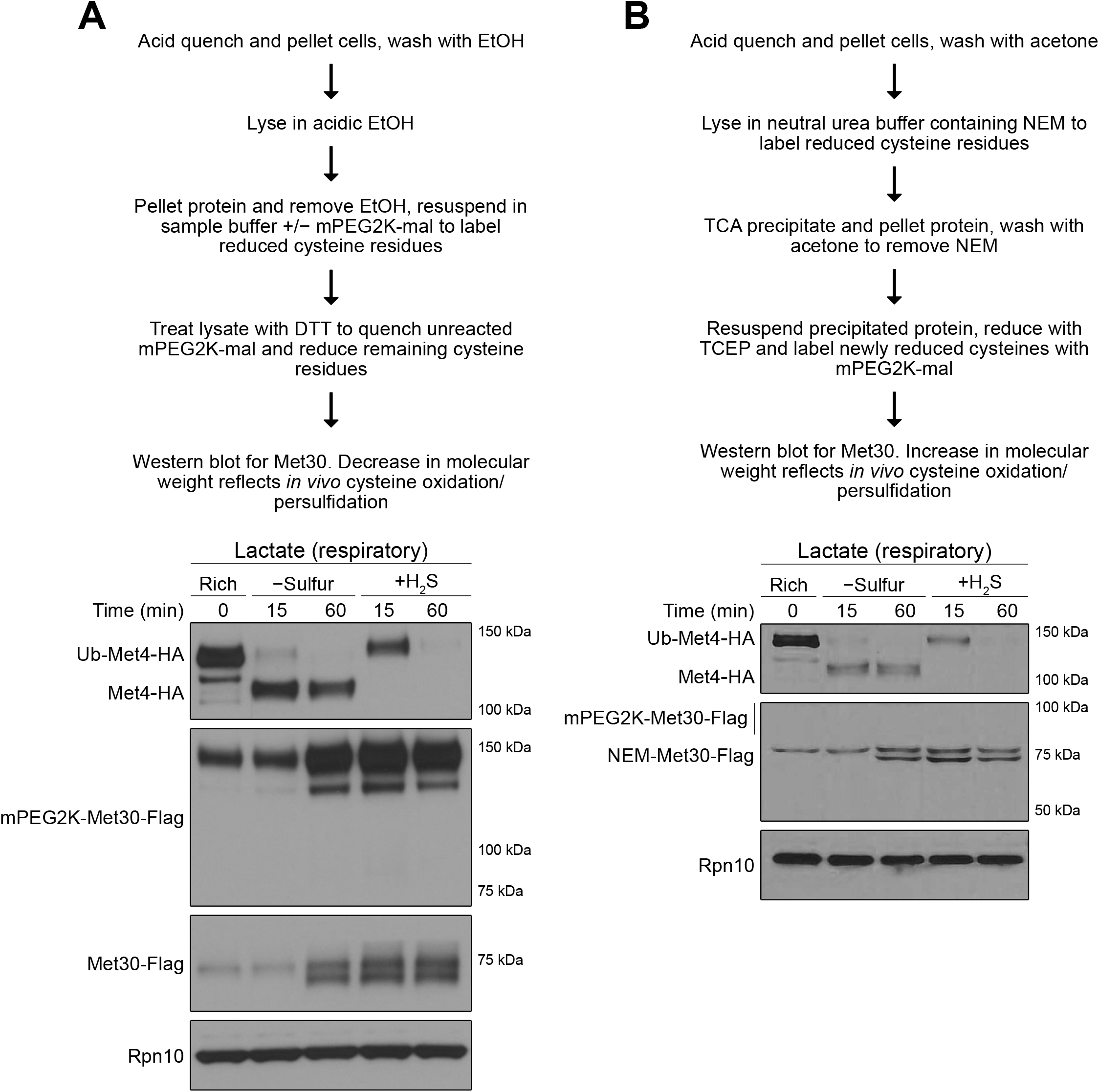
Cysteine labeling methods to detect Met30 cysteine oxidation or persulfidation. (A) Western blot of Met30 over the time course used in Figure 3. Cells were rapidly quenched in acid and lysed in acidic ethanol to preserve the redox state of Met30 cysteine residues. Precipitated protein was resuspended in sample buffer with or without mPEG2K-mal to label reduced cysteine residues in Met30 before reduction with DTT. Unmodified sample was used to blot for Met30 and Rpn10 as controls. (B) Western blot of Met30 was performed using a modified tag-switch method to detect changes in the redox or persulfidated status of Met30 cysteine residues over the sulfur starvation time course used in Figure 3 by mass shift. Cells were lysed in the presence of the small alkylating reagent NEM to modify all reduced or persulfidated protein thiols in the lysate, followed by acid precipitation of proteins, removal of residual NEM, resuspension and reduction by TCEP, and then a final alkylation step with mPEG2K-mal. The mPEG2K-mal reagent is expected to add approximately 2 kDa in mass for every oxidized (RSSR, RSOH, etc.) or persulfidated (RSSH) cysteine residue.

**Table S1.** Strains used in this study.

| BACKGROUND | GENOTYPE | SOURCE |
| --- | --- | --- |
| CEN.PK | MATa | (44) |
| CEN.PK | MATα | (44) |
| CEN.PK | MATa; MET30-FLAG::KanMX | This study |
| CEN.PK | MATa; MET30-FLAG::KanMX MET4-HA::Hyg | This study |
| CEN.PK | MATa; MET30-FLAG::KanMX MET4-HA::Hyg<br>met6Δ::Nat | This study |
| CEN.PK | MATa; MET30-FLAG::KanMX MET4-HA::Hyg<br>str3Δ::Nat | This study |
| CEN.PK | MATa; met30::MET30-C414S-FLAG::KanMX<br>MET4-HA::Hyg | This study |
| CEN.PK | MATa; met30::MET30-C614/616/622/630S-FLAG::KanMX MET4-HA::Hyg | This study |
| CEN.PK | MATa; met30Δ::Phleo HO::MET30-FLAG::Nat<br>MET4-HA::Hyg | This study |
| CEN.PK | MATa; met30Δ::Phleo HO::MET30Δaa1-20-FLAG::Nat Met4-HA::Hyg | This study |
| CEN.PK | MATa; met30Δ::Phleo HO::MET30-M30/35/36A-FLAG::Nat Met4-HA::Hyg | This study |
| CEN.PK | MATa; MET30-FLAG::KanMX MET4-HA::Hyg<br>pdr5Δ::Nat | This study |
| CEN.PK | MATa; met4Δ::KanMX MET30-FLAG::Hyg | This study |
| CEN.PK | MATa; met30::MET30-C201S-FLAG::KanMX<br>MET4-HA::Hyg | This study |
| CEN.PK | MATa; met30::MET30-C374S-FLAG::KanMX<br>MET4-HA::Hyg | This study |
| CEN.PK | MATa; met30::MET30-C426S-FLAG::KanMX<br>MET4-HA::Hyg | This study |
| CEN.PK | MATa; met30::MET30-C436S-FLAG::KanMX<br>MET4-HA::Hyg | This study |
| CEN.PK | MATa; met30::MET30-C439S-FLAG::KanMX<br>MET4-HA::Hyg | This study |
| CEN.PK | MATa; met30::MET30-C455S-FLAG::KanMX<br>MET4-HA::Hyg | This study |
| CEN.PK | MATa; met30::MET30-C528S-FLAG::KanMX<br>MET4-HA::Hyg | This study |
| CEN.PK | MATa; met30::MET30-C544S-FLAG::KanMX<br>MET4-HA::Hyg | This study |
| CEN.PK | MATa; met30::MET30-C584S-FLAG::KanMX<br>MET4-HA::Hyg | This study |
| CEN.PK | MATa; met30::MET30-C614S-FLAG::KanMX<br>MET4-HA::Hyg | This study |
| CEN.PK | MATa; met30::MET30-C616S-FLAG::KanMX<br>MET4-HA::Hyg | This study |
| CEN.PK | MATa; met30::MET30-C584/622S-FLAG::KanMX<br>MET4-HA::Hyg | This study |
| CEN.PK | MATa; met30::MET30-C630S-FLAG::KanMX<br>MET4-HA::Hyg | This study |
| CEN.PK | MATa; MET30-FLAG::KanMX MET4-HA::Hyg<br>met17Δ::Nat met10Δ::Phleo | This study |
| S288C | MATa; MET30-FLAG::KanMX MET4-HA::Hyg | This study |
| S288C | MATa; MET30-FLAG::KanMX MET4-HA::Hyg<br>cys3Δ::Nat | This study |
| S288C | MATa; MET30-FLAG::KanMX MET4-HA::Hyg<br>cys4Δ::Nat | This study |

**Table S2.**
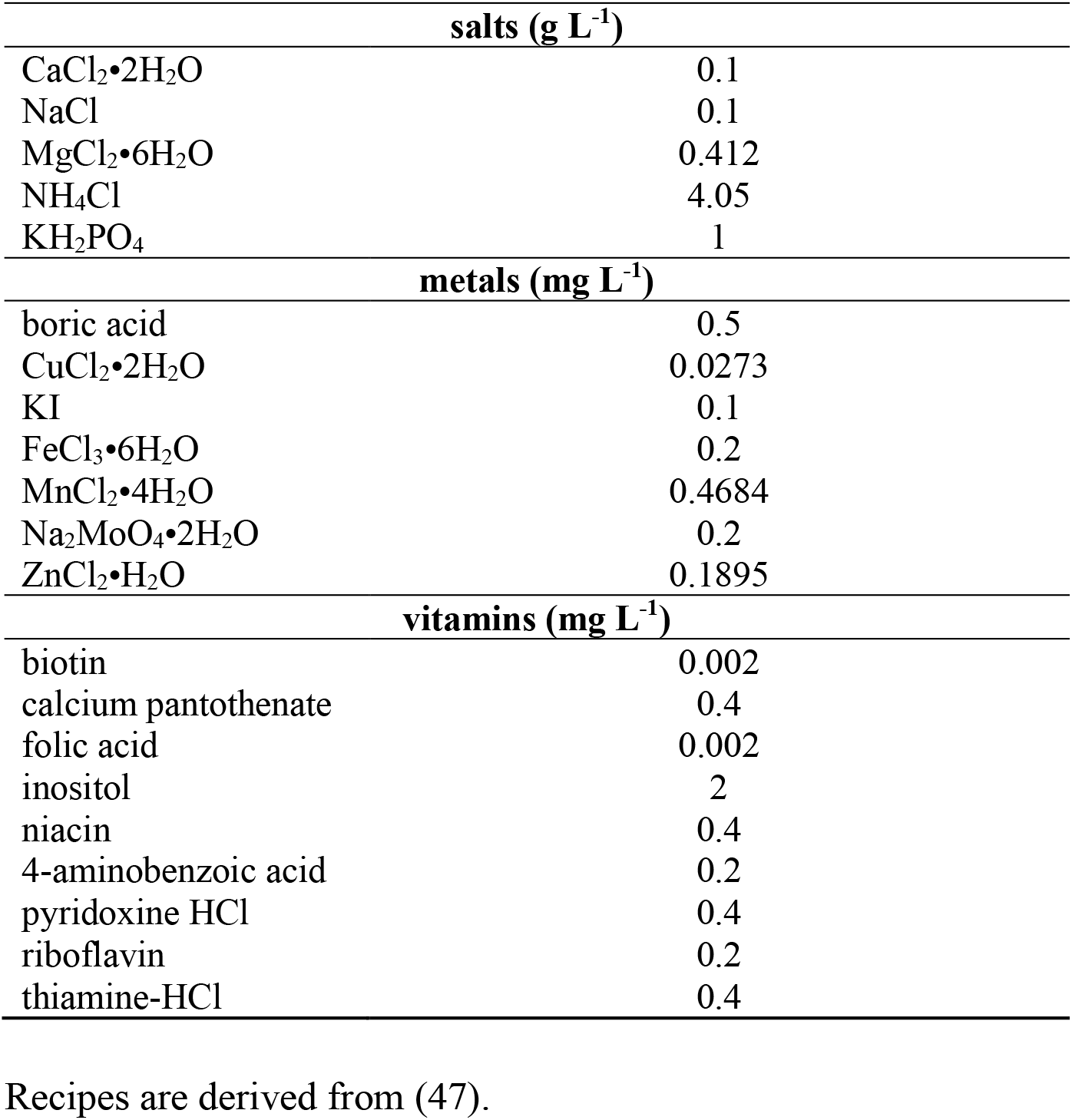
Recipe for sulfur-free media.

| **salts (g L<sup>-1</sup>)** |  |
| --- | --- |
| CaCl<sub>2</sub>•2H<sub>2</sub>O | 0.1 |
| NaCl | 0.1 |
| MgCl<sub>2</sub>•6H<sub>2</sub>O | 0.412 |
| NH<sub>4</sub>Cl | 4.05 |
| KH<sub>2</sub>PO<sub>4</sub> | 1 |
| **metals (mg L<sup>-1</sup>)** |  |
| boric acid | 0.5 |
| CuCl<sub>2</sub>•2H<sub>2</sub>O | 0.0273 |
| KI | 0.1 |
| FeCl<sub>3</sub>•6H<sub>2</sub>O | 0.2 |
| MnCl<sub>2</sub>•4H<sub>2</sub>O | 0.4684 |
| Na<sub>2</sub>MoO<sub>4</sub>•2H<sub>2</sub>O | 0.2 |
| ZnCl<sub>2</sub>•H<sub>2</sub>O | 0.1895 |
| **vitamins (mg L<sup>-1</sup>)** |  |
| biotin | 0.002 |
| calcium pantothenate | 0.4 |
| folic acid | 0.002 |
| inositol | 2 |
| niacin | 0.4 |
| 4-aminobenzoic acid | 0.2 |
| pyridoxine HCl | 0.4 |
| riboflavin | 0.2 |
| thiamine-HCl | 0.4 |

Recipes are derived from (47).

